# Photoinactivation of Catalase Sensitizes Wide-Ranging Bacteria to ROS-Producing Agents and Immune Cells

**DOI:** 10.1101/2021.06.24.449847

**Authors:** Pu-Ting Dong, Sebastian Jusuf, Jie Hui, Yuewei Zhan, Yifan Zhu, George Y. Liu, Ji-Xin Cheng

**Author notes:** These authors contributed equally: Pu-Ting Dong, Sebastian Jusuf, Jie Hui, Yuewei Zhan.

## Abstract

Bacteria have evolved to cope with the detrimental effects of reactive oxygen species (ROS) using their essential molecular components. Catalase, a heme-containing tetramer protein expressed universally in most of the aerobic bacteria, plays an indispensable role in scavenging excess hydrogen peroxide (H_2_O_2_). Here, through utilization of wild-type and catalase-deficient mutants, we identified catalase as an endogenous therapeutic target of 400-420 nm blue light. Catalase residing in bacteria could be effectively inactivated by blue light, subsequently rendering the pathogens extremely vulnerable to H_2_O_2_ and H_2_O_2_-producing agents. As a result, photoinactivation of catalase and H_2_O_2_ synergistically eliminate a wide range of catalase-positive planktonic bacteria and *P. aeruginosa* inside biofilms. In addition, photoinactivation of catalase is shown to facilitate macrophages to defend against intracellular pathogens. The antimicrobial efficacy of catalase photoinactivation is further validated using a *Pseudomonas aeruginosa-*induced mice abrasion model. Taken together, our findings offer a catalase-targeting phototherapy against multidrug-resistant bacterial infections.

## Introduction

Antibiotic resistance remains one of the biggest threats to global health over the past decades. In United States alone, it is estimated that at least 2.8 million people are infected by antibiotic-resistant bacterial infections annually^1^. Despite these alarming numbers, the pace of antibiotic resistance development is faster than that of clinical introduction of new antibiotics^2^. If no efforts are made to curtail this situation, life loss from antibiotic-resistant infections might surpass cancer, and 10 million people will be killed worldwide by 2050^3^. Moreover, the imprudent use of antibiotics from clinical over-prescription and food industry escalates the selection of multi-drug resistant or even pan-drug resistant bacteria^4^.

Confronted with this dire situation, antimicrobial blue light has emerged as a novel approach to combat multidrug-resistant bacterial infections^5,6^. Blue light in the 405-420 nm or the 450-470 nm optical windows has demonstrated bactericidal effects towards a wide range of microbial species, including gram-positive bacteria, gram-negative bacteria, mycobacteria, and molds^6^. Of note, blue light has been utilized for clinical treatment of *Propionibacterium acnes*^7^. *Helicobacter pylori*, the major cause of peptic ulcer disease, could be efficiently inactivated *in vitro* by visible light^8,9^. Besides planktonic-form bacteria, blue light also decreased the viability of *Pseudomonas aeruginosa* (*P. aeruginosa*), MRSA USA300 and *Candida albicans* in biofilm conditions^10^. Importantly, no evidence of blue light-resistance development by pathogens has been documented after consecutive blue light treatments^11-13^. Blue light has also been allied with other antimicrobial agents to eradicate bacteria. For example, quinine in combination with blue light exposure has shown efficacy to eliminate gram-negative *P. aeruginosa, Acinetobacter baumannii* (*A. baumannii*)^5^ and *Candida albicans*^14^. Blue light irradiance was also reported to enhance the inactivation efficacy of low-concentration chlorinated disinfectants towards *Clostridium difficile*^15^. Blue light (460 nm) plus hydrogen peroxide (H_2_O_2_) exhibited high efficacy to eradicate MRSA by devastating its functional membrane domain^16,17^.

Despite these advances, the working mechanism of antimicrobial blue light has remained elusive for years. Endogenous metal-free porphyrin or riboflavin has been considered as the major molecular targets^6^. It is assumed that reactive oxygen species (ROS) produced from photodynamic reaction between blue light and these endogenous chromophores lead to bacterial death. However, this hypothesis has remained controversial. The concentration of endogenous porphyrins or riboflavin is as low as 2-4×10^−3^ mg/ml^18^, and a precursor, δ-aminolevulinic acid (ALA), was routinely administered to enhance the intracellular production of porphyrins when treating *E. coli* and other bacteria^19^ by 400-420 nm blue light. In the absence of ALA, 407-420 nm blue light with a dose of 50 J/cm^2^ did not exhibit significant bactericidal effects on *Staphylococcus aureus* (*S. aureus*), *A. baumannii* and *E. coli*^19^. Also, it has been reported that the total amount of coproporphyrins was not a contributing factor of the antimicrobial efficacy of blue light treatment^20^. Alternatively, pyocyanin, the prototypical green pigment produced by *P. aeruginosa*^21^, has been suggested to serve as a photosensitizer upon blue light exposure^22,23^. Pyoverdine, a naturally occurring fluorescent pigment in *P. aeruginosa*, was also believed to undergo photodynamic reactions upon absorption of 415 nm light^24^.

Very recently, our studies found that blue light at 460 nm is able to bleach staphyloxanthin^16^, a ROS scavenger as well as an endogenous golden pigment residing in *S. aureus* functional membrane domains^25,26^, makes this pathogen vulnerable to low-concentration H_2_ O_2_ ^16^. Follow-up studies using pulsed blue light have shown more effective capability of photobleaching of staphyloxanthin, which sensitizes *S. aureus* to a broad spectrum of antibiotics^17^ and to silver nanoparticles^27^. In an independent study, it was shown that photobleaching of another ROS scavenger and pigment, granadaene, by 430 nm light is able to reduce the virulence and increase the antimicrobial susceptibility of *Streptococcus agalactiae*^28^. Collectively, these findings suggest an alternative working mechanism of antimicrobial blue light, which is based on photo-inactivation of intrinsic ROS-scavenging molecules inside the pathogen.

It is well established that aerobic microorganisms produce ROS endogenously when flavin, quinol, or iron cofactors are autoxidized in the process of cellular metabolism and respiration^29^. When bacteria are challenged with antibiotics or other stressors, a cascade of ROS could be generated^30,31^. Excess ROS damage DNA, certain metalloproteins, lipids, and other essential cellular components^32^. To scavenge the excess ROS and maintain the intracellular homeostasis, bacteria have evolved to be armed with array of strategies. Of these, catalase, an enzyme with a turnover number of 2.8×10^6^ molecules per second^33^, very efficiently converts H_2_O_2_ into O_2_ and water. For this reason, visualization of oxygen bubbles in the presence of H_2_O_2_ and Triton-X offered a simple method to quantify catalase activity^34^. In the absence of catalase, Fenton reaction between H_2_O_2_ and iron would produce various radicals, such as HO? and HOO?, and poses lethal threats to bacteria^35^. Importantly, it was shown as early as 1965 that catalase could be inactivated by visible light^36^, with the optical density at 405 nm (the primary absorption peak) diminishing as the irradiance continued. The underlying mechanism was presumed to be due to the dissociation of prosthetic heme groups from the tetramer protein^37^. Nonetheless, whether photoinactivation of catalase could be harnessed to eliminate pathogenic bacteria is yet to be explored.

Here, we show that catalase expressed universally in common pathogens is a key target of anti-microbial blue light in the 400-420 nm optical window. Blue light illumination inactivates catalase by destroying the porphyrin rings. Using the same dose, nanosecond (ns) pulsed blue light at 410 nm induced more effective inactivation of catalase than the continuous-wave (CW) 410 nm irradiance. We further demonstrate that photoinactivation of catalase sensitized bacterial pathogens, both in planktonic form and biofilms, to exogenous non-bactericidal low-concentration H_2_O_2_. Moreover, photoinactivation of catalase sensitized pathogens to certain antibiotics that exert their lethal effects on bacteria partly through ROS-induced physiological alterations^38^. Photoinactivation of catalase enhanced immune cell elimination of intracellular MRSA USA300 and *P. aeruginosa*. In a *P. aeruginosa*-induced mouse skin abrasion model, photoinactivation of catalase effectively reduced the pathogen burden. Taken together, our findings offer a new way to combat multi-drug resistant bacterial infections.

## Results

### Bacterial catalase can be inactivated by blue light irradiance

Catalase is a tetramer of 60,000-dalton subunits, containing four prosthetic heme rings per tetramer. As shown in **Figure 1a**, the porphyrin rings stay deep inside the hydrophobic pocket of catalase from *P. aeruginosa*. Since bovine liver catalase has been shown to be inactivated by visible light decades ago^36^ but with unclear underlying mechanism, we asked whether and how catalase structure changes in the presence of light exposure near its absorption around 405 nm. For this purpose, Raman spectra of a dried bovine liver catalase film were obtained before and after 410 nm exposure. Catalase exhibits its prototypical peaks at 754 cm^-1^ and 1200 cm^-1^ to 1500 cm^-1^ due to porphyrin rings (**Figure 1b**)^39^. Interestingly, the Raman intensity at those peaks drastically dropped after 410 nm exposure, indicating a possible dissociation of the heme ring from catalase.

**Figure 1.**
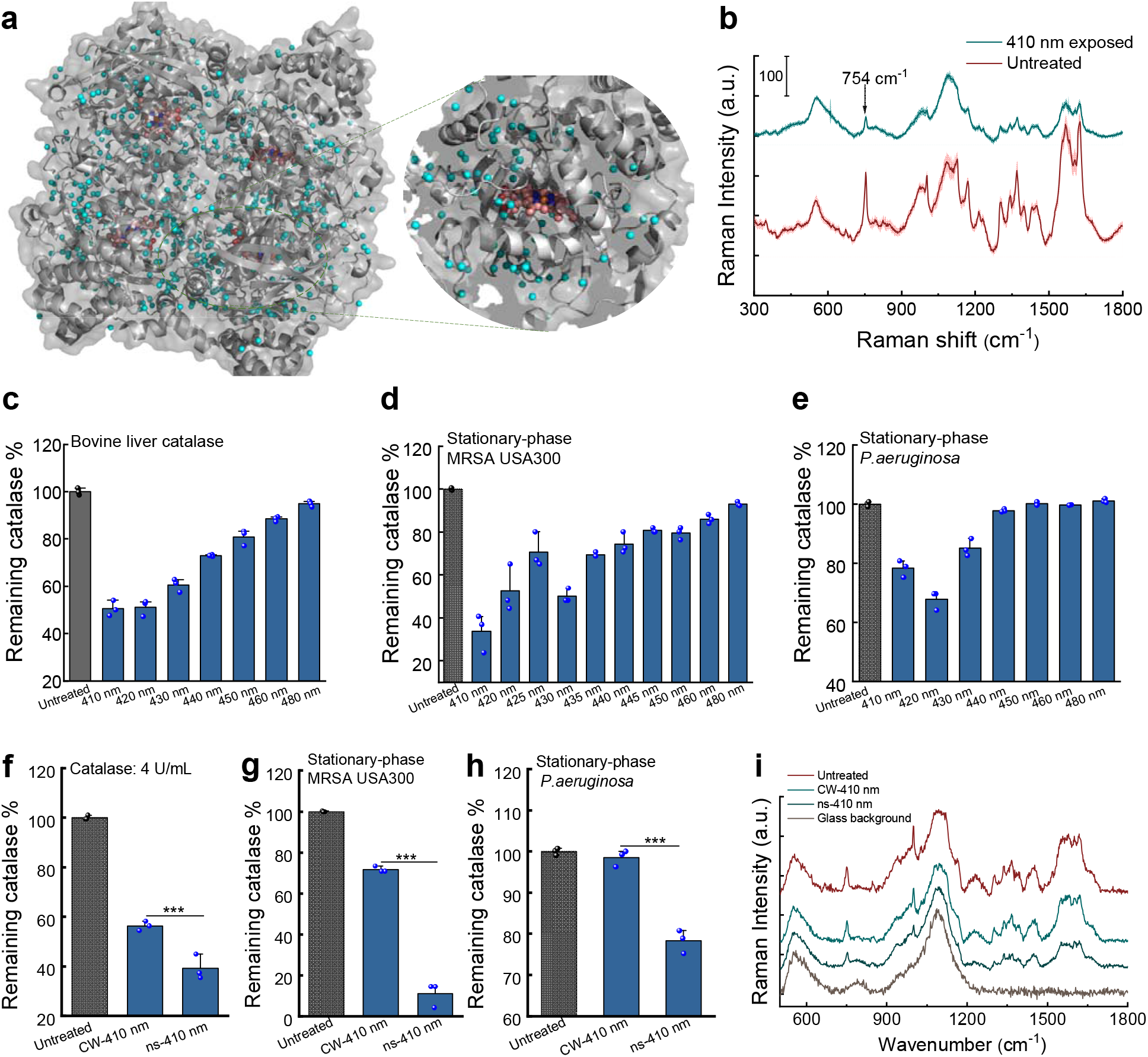
Catalase inactivation by CW and pulsed blue light in a wavelength dependent manner. **a**. *P. aeruginosa* catalase structure through PyMOL simulation. PDB 4E37. **b**. Raman spectra of a bovine liver catalase film dried on a cover slide before (untreated) and after 410 nm light exposure. The Raman peak at 754 cm^-1^ was highlighted by an arrow. 410 nm: 50 mW/cm^2^, 10 min. Standard deviations at each Raman shift were shown in shadow. **c**-**e**. Remaining catalase percent of bovine liver catalase (**c**, 2.5 U/ml), stationary-phase MRSA USA300 (**d**) and stationary-phase *P. aeruginosa* PAO1 (**e**) under blue light exposure at different wavelengths. **f**-**h**. Comparison of continuous-wave (CW) and ns-410 nm exposure on inhibiting bovine liver catalase (**f**, 4 U/ml), catalase from stationary-phase MRSA USA300 (**g**) and stationary-phase *P. aeruginosa* PAO1 (**h**). **i**. Raman spectra of bovine liver catalase film dried on a cover slide under CW-410 and ns-410 exposure. Light: 50 mW/cm^2^, 5 min. Data: Mean±SD. Statistical analysis was obtained through student unpaired *t*-test. ***: *p*<0.001.

To further understand the impact of 410 nm exposure on catalase, transient absorption microscopy was utilized to record the time-resolved photobleaching behavior of bovine liver catalase, and catalase inside MRSA USA300, *P. aeruginosa*, and *Salmonella enterica* (**supplementary Figure 1**). Samples were excited by a pump beam at wavelength of 410 nm and transient absorption signals were detected by a probe beam at wavelength of 520 nm. As shown in **supplementary Figure 1**, the transient absorption signal from bovine liver catalase decayed as the irradiance continued (**supplementary Figure 1, a-c**). Moreover, this decay (**supplementary Figure 1d)** follows a second-order photobleaching model^16^, suggesting an interaction between heme rings within the tetramer. Consistently, photoinactivation of catalase from MRSA USA300 (**supplementary Figure 1e**), *P. aeruginosa* (**supplementary Figure 1f**), and *Salmonella enterica* (**supplementary Figure 1g**) followed the similar photobleaching trend as that of bovine liver catalase. Functionally, after 410 nm exposure, no apparent bubbles were observed when adding 0.3% H_2_O_2_ to the light-exposed catalase solution (**supplementary Video 1**). In summary, the enzyme-bound heme rings could be dissociated from the protein upon 410 nm irradiance, causing the protein to malfunction.

Next, we asked what is the optimal wavelength to inactivate catalase. To investigate this, we employed an OPO laser (Opolette 355, OPOTEK) and varied the laser wavelength under the same power density and dose. A catalase kit that quantifies the residual H_2_O_2_ was then used to indirectly measure the remaining catalase percent in order to evaluate the efficacy of light exposure. As shown in **Figure 1c**, 410-420 nm demonstrated the highest efficiency to inactivate bovine liver catalase (∼50% inactivation at a dose of 15 J/cm^2^). Catalase plays a dominant role in conversion of H_2_O_2_ into water and O_2_ in bacteria^40^. We adopted the same protocol to evaluate the remaining catalase percentage in stationary-phase MRSA USA300 (**Figure 1d**) and *P. aeruginosa* (**Figure 1e**). Consistently, 410-420 nm exposure most effectively attenuated the bacterial catalase activity. Therefore, 410 nm was utilized for catalase inactivation for the subsequent experiments.

Since photoinactivation of catalase is likely due to the dissociation of the heme rings from the protein following a second-order photobleaching model, we asked whether high-peak power pulsed (e.g. nanosecond) 410 nm (ns-410) blue light more effectively inactivates catalase compared to the 410 nm continuous wave (CW-410). To answer this question, three experiments were conducted. First, we used the same catalase kit to quantify the H_2_O_2_ conversion efficacy of bovine liver catalase (**Figure 1f**), stationary-phase MRSA USA300 (**Figure 1g**), and stationary-phase *P. aeruginosa* (**Figure 1h**) after ns-410 and CW-410 blue light exposure, respectively. Our data repeatedly exhibited the pattern that ns-410 is significantly more effective for catalase inactivation compared to CW-410 (*p*<0.001). Second, Raman spectra of bovine liver catalase film dried on a glass substrate were obtained after ns-410 and CW-410 exposure. Noteworthy, under the same dose, ns-410 nm displayed a better capability to bleach catalase as Raman peaks at 754 cm^-1^ and 1200 to 1500 cm^-1^ drastically dropped (**Figure 1i**). Third, we compared the heights of oxygen bubble foams produced by catalase in the presence of H_2_O_2_ and Triton-X after ns-410 and CW-410 exposure at the same dose. As shown in **supplementary Figure 2**, ns-410 treated groups indeed showed much lower foam heights. Collectively, ns-410 exposure is more effective to inactivate catalase compared to CW-410. Notably, ns-410 exposure also alleviates the heat accumulation issue which generally concomitants with long-time CW light exposure.

### Photoinactivation of catalase sensitizes pathogenic bacteria to low H_2_O_2_ -concentrations

Since the primary role of catalase is to degrade H_2_O_2_ into water and oxygen, we then investigated whether photoinactivation of catalase renders bacteria vulnerable to H_2_O_2_. We tested the viability of MRSA USA300 and *P. aeruginosa* after different treatments (various light wavelength, light dose, and H_2_O_2_ concentration) by colony-forming units (CFU/ml) enumeration. H_2_O_2_ at 22 mM (or 0.075%) didn’t show apparent bactericidal effect (**Figure 2, a-b**). Yet, 4-log_10_ reduction was achieved in the 410 nm plus H_2_O_2_ treated group, indicating a very strong synergy between 410 nm exposure and H_2_O_2_ for bacterial eradication.

**Figure 2.**
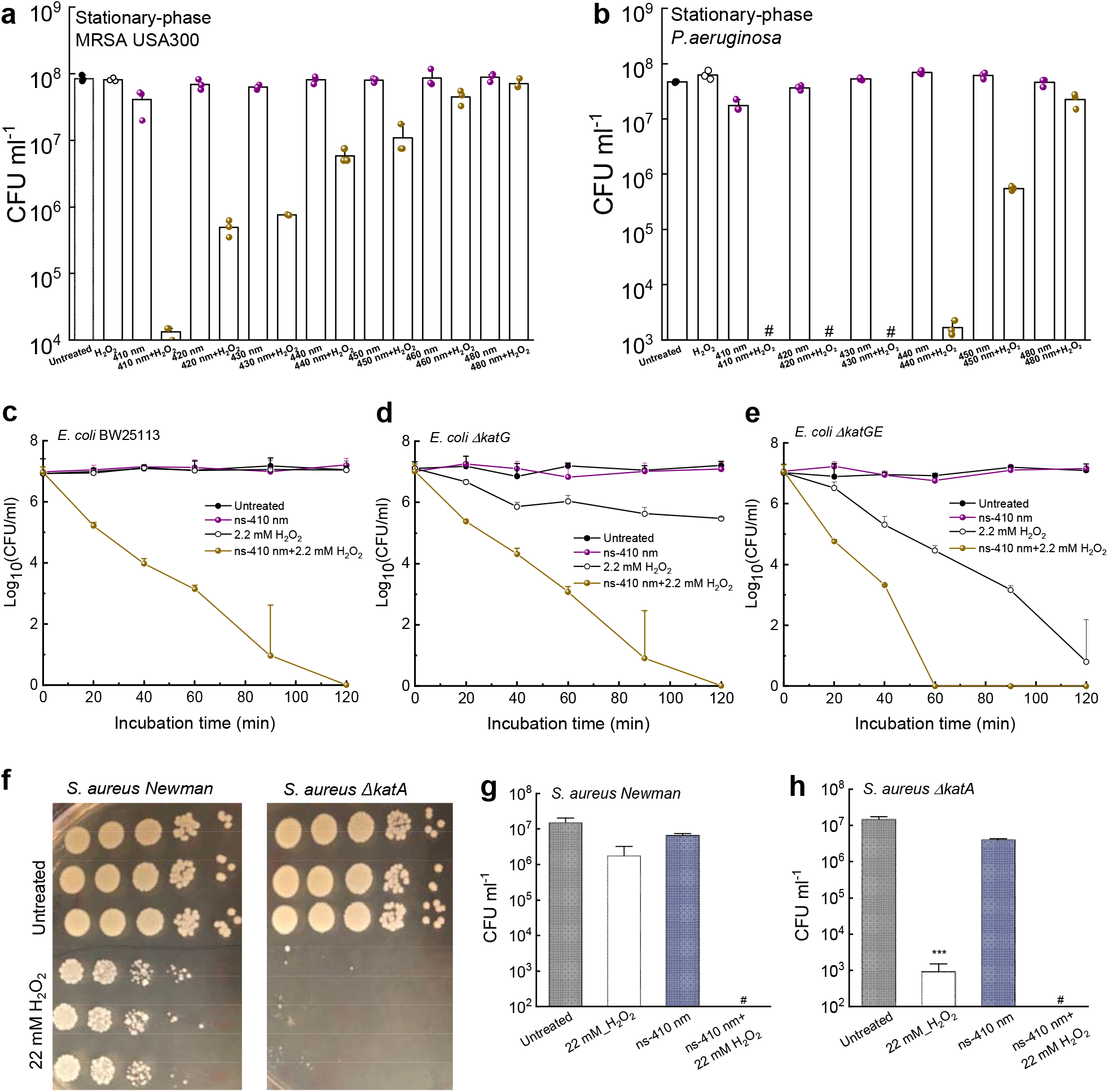
Photoinactivation of catalase effectively sensitizes pathogenic bacteria to H_2_O_2_. **a**. CFU ml^-1^ of stationary-phase MRSA USA300 under various treatments. Dose: 50 mW/cm^2^, 5 min. H_2_O_2_: 22 mM, 30-min incubation time. **b**. CFU ml^-1^ of stationary-phase *P. aeruginosa* PAO1 under various treatments. Dose: 50 mW/cm^2^, 5 min. H_2_O_2_: 22 mM, 30-min incubation time. **c-e**. Time-killing curves of wild type *E. coli* BW25113 (**c**), *E. coli ΔkatG* (**d**), *E. coli ΔkatGE* (**e**) under different treatment schemes. ns-410 nm: 30 mW, 8 min, 14 J/cm^2^. **f**. CFU plates of *S. aureus Newman* along with its isogenic catalase mutant *S. aureus ΔkatA* with/without H_2_O_2_ treatment. **g-h**. CFU ml^-1^ of *S. aureus Newman* (**g**) and *S. aureus ΔkatA* (**h**) under different treatment schemes. Data: Mean±SD. Pound sign (#) indicates the CFU results are below the detection limit. Student unpaired *t*-test (compared to the untreated group). ***: *p*<0.001.

It is noteworthy that the more recalcitrant stationary-phase *P. aeruginosa* PAO1 was completely eradicated by 410 nm plus H_2_O_2_ (**Figure 2b**). When tuning the irradiance wavelengths, the killing pattern is similar to that of catalase inactivation (**Figure 2, a-b**), suggesting catalase is a key target of blue light irradiance. In addition, ns-410 plus H_2_O_2_ treatment of MRSA USA300 outperformed CW-410 plus H_2_O_2_ treatment by approximately 3-log_10_ (**supplementary Figure 3a**). The combination treatment that made use of ns-410 achieved total eradication of *P. aeruginosa* PAO1, and outperformed the CW-410 plus H_2_O_2_ treatment (**supplementary Figure 3b**). These findings are in line with ns-410 being more efficient at inactivating catalase than CW-410 when using the same dose.

To further investigate whether it is catalase that primarily accounts for the synergy between 410 nm exposure and H_2_O_2_, we measured the viability of *E. coli* BW25113 along with its catalase-deficient mutants, *E. coli ΔkatG* (single mutant) and *E. coli ΔkatGE* (double mutant) under the same treatments. First, a time-killing CFU assay was conducted. Complete eradication of *E. coli* BW25113 was obtained in the ns-410 plus H_2_O_2_ treated group after two hours of incubation in PBS whereas 410 nm alone or H_2_O_2_ alone barely killed the bacteria (**Figure 2c**). The synergistic activity of a combination treatment is defined as a >2-log_10_ decrease in the CFU ml^-1^ compared to that obtained with the most active agent alone^41^. Therefore, there is an effective synergy between 410 nm blue light and H_2_O_2_.

Catalase-deficient *E. coli ΔkatG* and *E. coli ΔkatGE* have baseline susceptibility to H_2_O_2_ that are drastically higher compared to the isogenic wild type *E. coli* (**Figure 2, d-e**). The double mutant *E. coli ΔkatGE* exhibited no visible oxygen bubbles formation with the addition of H_2_O_2_ (**supplementary Figure 4**). Moreover, *E. coli ΔkatGE* exhibited similar susceptibility to H_2_O_2_ killing compared to wild type *E. coli* BW25113 exposed to ns-410 plus H_2_O_2_ in a time-kill assay, corroborating that catalase is the primary target of 410 nm light. More effective killing of the catalase-deficient mutants by 410 nm plus H_2_O_2_ was observed when compared to H_2_O_2_ treatment alone, suggesting the existence of additional molecular targets for 410 nm exposure.

To further confirm that catalase is the primary target of blue light and inactivation of its underlying function accounts for the synergy between 410 nm and H_2_O_2_, we further applied the treatments to *S. aureus Newman* along with its isogenic catalase-deficient mutant *S. aureus ΔkatA*. At a concentration of 22 mM, H_2_O_2_ eliminated less than 1-log_10_ of *S. aureus Newman* yet reduced around 4-log_10_ of *S. aureus ΔkatA* (**Figure 2f**). When combining 410 nm exposure and H_2_O_2_, total eradications were obtained for both *S. aureus Newman* (**Figure 2g**) and *S. aureus ΔkatA* (**Figure 2h**). These data further suggest that catalase is the primary molecular target for 410-nm light exposure.

### Catalase photoinactivation and H_2_O_2_ synergistically eradicate wide-ranging bacteria

Having established the synergy between 410 nm exposure and H_2_O_2_ in eradication of *E. coli*, MRSA USA300, and *P. aeruginosa*, we next asked whether this synergy works on other pathogenic bacteria. In 2017, the world health organization published a list of twelve global priority pathogens, including *Acinetobacter baumannii* (*A. baumannii*), *P. aeruginosa, Salmonellae* etc^42^, most of which are catalase positive. It was reported that *A. baumannii* could induce severe mucous membrane infections or even bacteremia^43^, *P. aeruginosa* has been the main culprit among patients with burn wounds, cystic fibrosis, acute leukemia, organ transplants^44^. *Salmonellae*, the most commonly isolated foodborne pathogens, lead to approximately three million deaths worldwide due to *Salmonella* gastroenteritis^45^. Therefore, we explored whether 410 nm exposure and H_2_O_2_ are effective against these life-threatening pathogens.

Consistently, we found that total eradications (around 8-log_10_ reduction) were achieved when the treatment was applied to stationary-phase *Salmonella enterica* ATCC 29630 (**Figure 3a**) and stationary-phase *E. coli* BW25113 (**Figure 3b**). 5-log_10_ reduction of stationary-phase *A. baumannii 1* (**Figure 3c**) was achieved in the 410 nm plus H_2_O_2_ treated group. Notably, this synergistic cocktail therapy achieved total eradication (around 8-log_10_ reduction) of multiple *P. aeruginosa strains* (**Figure 3, d-f**) and *Klebsiella pneumonia 1* strains (**Figure 3g**). We then asked if this synergy still holds for catalase-negative pathogens, e.g. *Enterococcus faecalis* (*E. faecalis 1*)^46^. As shown in **Figure 3h**, H_2_O_2_ alone effectively killed *E. faecalis 1* by around 3-log_10_, but the enhanced bactericidal effect associated with 410 nm exposure was not as significant as that for catalase-positive pathogens. Taken together, 410 nm plus H_2_O_2_ can effectively eradicate wide-ranging life-threatening pathogens.

**Figure 3.**
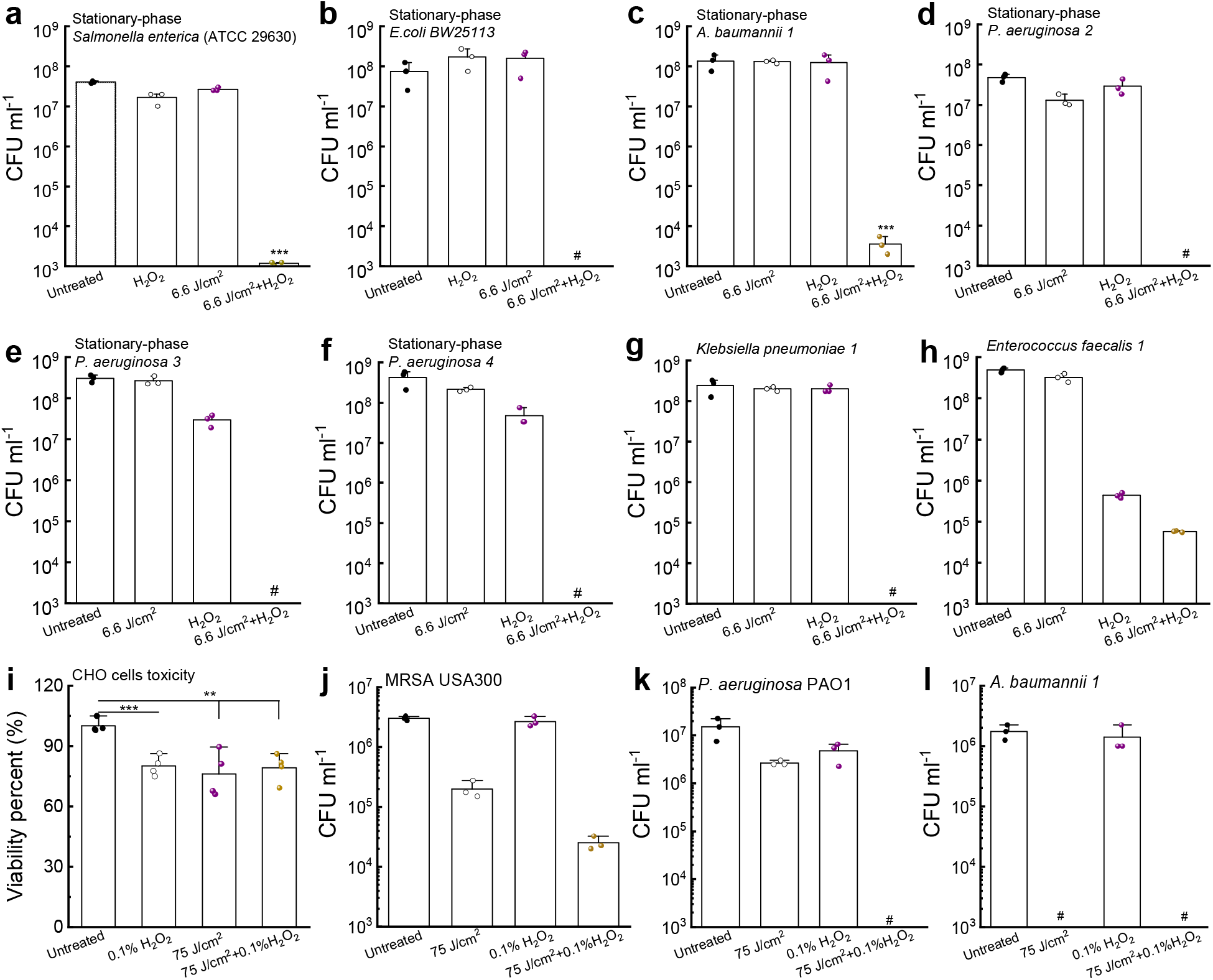
Photoinactivation of catalase sensitizes wide-ranging pathogenic bacteria to H_2_O_2_. **a**-**h**. CFU ml^-1^ of stationary-phase *Salmonella enterica* ATCC 29630 (**a**), stationary-phase *E. coli* BW25113 (**b**), stationary-phase *A. baumannii 1* (**c**), stationary-phase *P. aeruginosa* strains (**d**-**f**), *Klebsiella pneumonia 1* (**g**), *Enterococcus faecalis 1* (**h**) under various treatment schemes. 410 nm exposure: 6.6 J/cm^2^, H_2_O_2_: 22 mM, 30-min culture time (for **a**-**h**). **i**. Toxicity test of CHO cells by MTT assay under different treatment schemes. **j**-**l**. CFU ml^-1^ of log-phase MRSA USA300 (**j**), log-phase *P. aeruginosa* PAO1 (**k**), and log-phase *A. baumannii 1* (**l**) under different treatment schemes. 410 nm exposure: 75 J/cm^2^, H_2_O_2_: 0.1%. Serial dilution or MTT assay was conducted after 1-min incubation time (for **i**-**l)**. Data: Mean±SD. Pound sign (#) indicates the CFU results are below the detection limit. Student unpaired *t*-test. ***: *p*<0.001.

To query the translational potential of this synergistic therapy, we inquired whether short H_2_O_2_ (with higher concentration) incubation time remains effective to eliminate bacteria after 410 nm exposure. Prior to that, we evaluated the toxicity of 0.1% (w/v) H_2_O_2_ (30 mM, 2-min incubation), 410 nm exposure (75 J/cm^2^), 410 nm exposure plus 0.1% H_2_O_2_ to a Chinese Hamster Ovary (CHO) cell line through MTT assay. A 0.1%∼0.2% of H_2_O_2_ was chosen since it was reported that H_2_O_2_ in the commercially disinfectant formulations is in the range of 0.1% to 3%^47^. As shown in **Figure 3i**, viability of CHO cells (around 80%) treated with 410 nm (75 J/cm^2^) plus H_2_O_2_ (0.1%, 1-min incubation) was similar CHO cells treated with 410 nm or H_2_O_2_.

Hence, we moved next to interrogate the bactericidal effect of short treatment against log-phase MRSA USA30, *P. aeruginosa* PAO1 and *A. baumannii 1*. Enhanced killing of log-phase MRSA USA300 was observed with 410 nm plus H_2_O_2_ treatment compared to treatment with H_2_O_2_ or 410 nm alone (**Figure 3j**). Total eradication was achieved in the case of log-phase *P. aeruginosa* PAO1 (**Figure 3k**) and log-phase *A. baumannii 1* (**Figure 3l**) with 410 nm and 1-min 0.1%-0.2% H_2_O_2_. Log-phase MRSA USA300, *P. aeruginosa* PAO1 and *A. baumannii 1* were particularly sensitive to 410 nm exposure (**Figure 3l**) compared to the bacteria grown to stationary-phase, indicating possible variation in catalase expression with growth condition. Collectively, 410 nm plus short exposure to H_2_O_2_ are sufficient to achieve significant bacterial reduction with negligible toxicity.

### Photoinactivation of catalase enhances microbicidal activity of H_2_O_2_-producing antibiotics

Significant intracellular H_2_O_2_ is produced upon treatment with ampicillin (β-lactam), gentamicin (aminoglycoside), and norfloxacin (fluoroquinolone)^38^. Based on these prior studies, we reasoned that photoinactivation of catalase could enhance the efficacy of these antibiotics that induce ROS generation. To test our hypothesis, we firstly tested tobramycin, a representative of aminoglycoside that could indirectly enhance the intracellular production of H_2_O_2_^48^. Augmented killing effect was observed with 410 nm exposure plus tobramycin treatment of *E. coli* BW25113 compared with tobramycin treatment alone (**Figure 4a**). 410 nm exposure strengthened the bactericidal effect of tobramycin against *Salmonella enterica* by around 1-log_10_ (**Figure 4, b-c**), and did not enhance tobramycin killing of catalase-negative *Enterococcus faecalis* 1 (**Figure 4d**).

**Figure 4.**
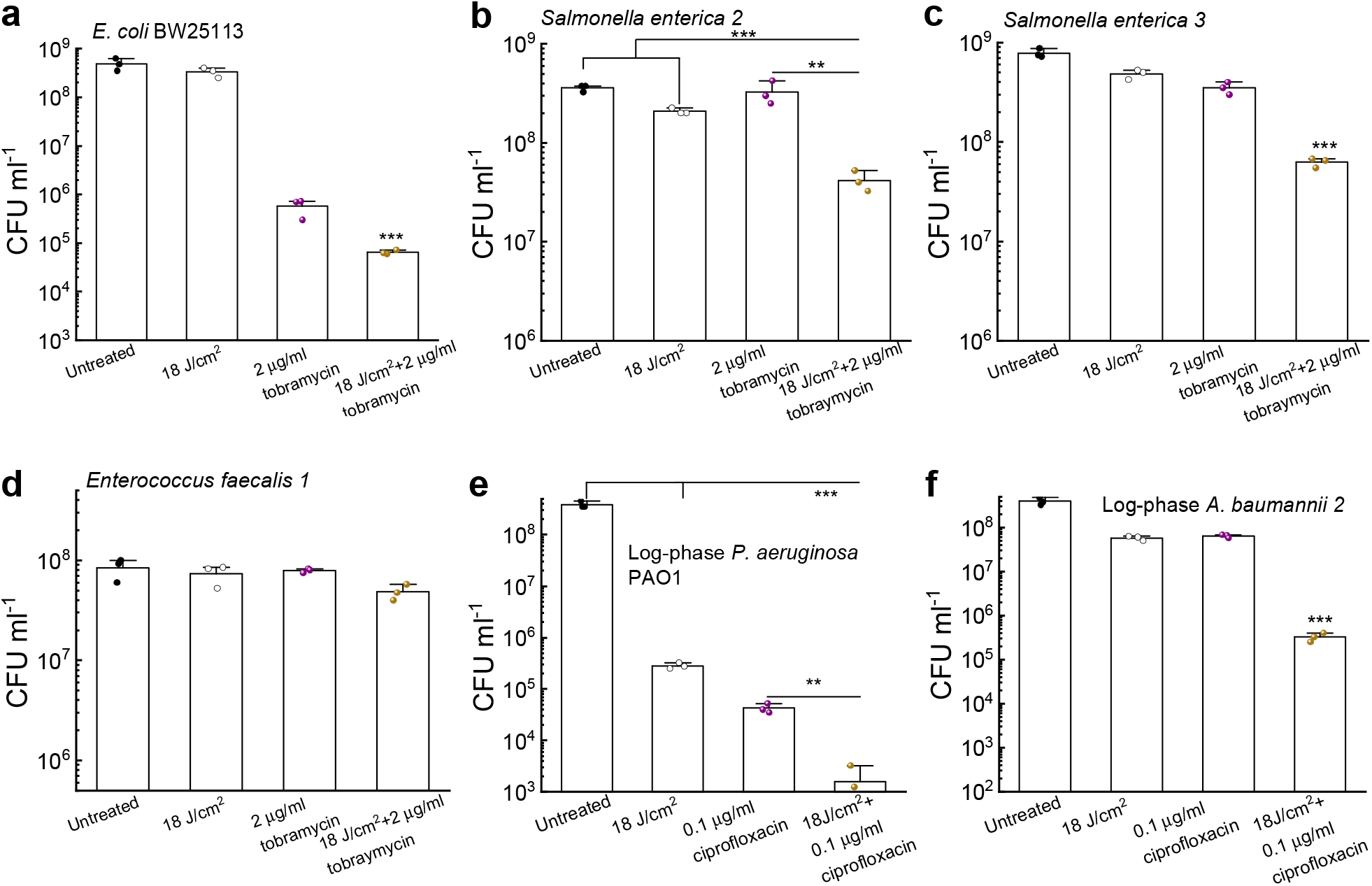
Photoinactivation of catalase sensitizes pathogenic bacteria to certain antibiotics. **a**. CFU ml^-1^ of *E. coli* BW25113 under different treatments. **b**-**c**. CFU ml^-1^ of *Salmonella enterica* under different treatments. CFU enumeration was obtained by incubation with tobramycin in TSB for 4 hours. **d**. CFU ml^-1^ of *Enterococcus faecalis 1* under the same treatments as in **b**-**c. e**-**f**. CFU ml^-1^ of log-phase *P. aeruginosa* PAO1 (**e**) along with *A. baumannii 2* under different treatment schemes. CFU enumeration was achieved after incubation with ciprofloxacin in TSB for 4 hours. Data: Mean±SD. Student unpaired *t*-test and one-way ANOVA. ***: *p*<0.001, **: *p*<0.01. Significant from other groups unless notified.

Per report, catalase-deficient *E. coli* mutants are more susceptible to fluoroquinolone ciprofloxacin when compared to their parent strain^49^. We thus wondered whether photoinactivation of catalase could sensitize bacteria to ciprofloxacin. As shown in **Figure 4e**, photoinactivation of catalase indeed boosted the antimicrobial potency of ciprofloxacin to eliminate log-phase *P. aeruginosa* PAO1. Similar enhancement was observed for log-phase *A. baumannii* 2 (**Figure 4f**). Collectively, these findings suggest that photoinactivation of catalase augments the antimicrobial efficacy of at least some antibiotics that indirectly increase the intracellular H_2_O_2_ level.

### Photoinactivation of catalase and H_2_O_2_ synergistically eliminate *P. aeruginosa* biofilms

Having shown effective synergy between photoinactivation of catalase and ROS-generating agents against both log- and stationary-phase planktonic bacteria, we next asked if the synergy can effectively eradicate biofilm-dwelling bacteria.

To address our query, we selected *P. aeruginosa* as a target as it is a notorious pathogen in burn wounds, chronic obstructive pulmonary disorder and cystic fibrosis^50^. Extensive *P. aeruginosa* biofilm protect the bacterium from host defense, chemotherapy and conventional antimicrobial therapy, leading to undesirable disease burden on patients^51,52^.

To mimic clinical *P. aeruginosa* infections in which biofilm has a major presence, we adopted a CDC biofilm growth protocol which grows *P. aeruginosa* biofilms on a polypropylene coupon under continuous flow conditions^53^. After forming robust biofilms, we applied treatments (untreated, H_2_O_2_ at a series of concentrations, 410 nm exposure (21 J/cm^2^), 410 nm (21 J/cm^2^) plus H_2_O_2_ (30-min incubation time)). Then we used a Live (SYTO 9)/Dead (Propidium iodide, PI) staining kit to image live/dead *P. aeruginosa* through a confocal microscope. As shown in **Figure 5a_1_-d_1_**, H_2_O_2_ treated groups barely have noticeable number of dead cells compared to the untreated group (**Figure 5e**). In comparison, drastic increase of dead bacteria number was observed in groups treated with 410 nm exposure plus H_2_O_2_ (**Figure 5, a_2_-d_2_, f**). 3D rendered images of *P. aeruginosa* biofilms (**supplementary Video 2**) further corroborated the synergistic effect between 410 nm exposure and H_2_O_2_ treatment to eliminate *P. aeruginosa* biofilms.

**Figure 5.**
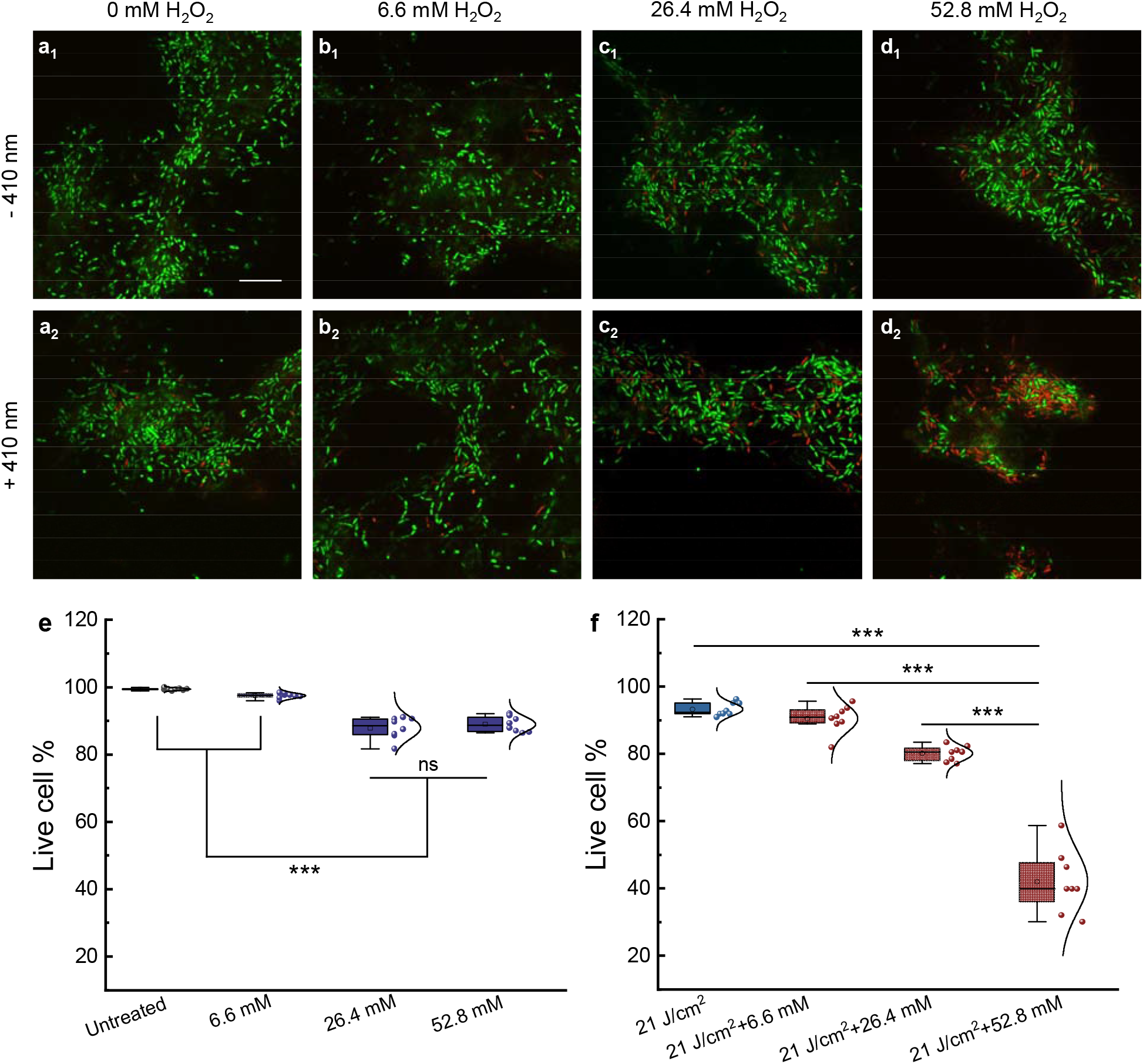
Confocal laser scanning microscopy of live/dead bacteria inside *P. aeruginosa* PAO1 biofilms after different treatments. **a**_1_-**d**_1_. Merged live/dead *P. aeruginosa* biofilms with various H_2_O_2_ treatments. **a**_**2**_**-d**_**2**_. Merged live/dead *P. aeruginosa* biofilms after 410 nm exposure combined with various H_2_O_2_ treatments. **e-f**. Quantitative analysis of the live cell percent of *P. aeruginosa* biofilms among the above eight groups. Data: Mean±SD from eight different field of views. Scalar bar=10 µm. Live: SYTO 9. Dead: PI. 410 nm laser: ns-410 nm, 35 mW, 10 min. H_2_O_2_: 30-min incubation time. Statistical analysis was determined through student unpaired *t*-test and one-way ANOVA. ***: *p*<0.001, ns: not significant.

Of note, even after combining 410 nm and H_2_O_2_ at a concentration of 52.8 mM, full eradication still could not be achieved (around 70% elimination, **Figure 5d_2_, f**), which may be attributed to the shielding conferred by the extracellular polymeric substances or quorum sensing inside the biofilms. Nonetheless, photoinactivation of catalase and H_2_O_2_ substantially eliminated *P. aeruginosa* in the biofilm setting, which suggests the potential for treating clinically relevant *P. aeruginosa* infections such as burn infections.

### Photoinactivation of catalase assists macrophages to eliminate intracellular pathogens

Besides forming persistent biofilms, bacteria could also reside inside host immune or non-immune cells to evade antibiotic attack^54,55^. It has been reported that the minimal inhibitory concentrations (MICs) of intracellular MRSA is two orders of magnitude higher than those of free-living planktonic bacteria^55^. Furthermore, *S. aureus* could proliferate within host phagocytic cells such as neutrophils and macrophages shortly after intravenous infection^56^. These viable intracellular *S. aureus* allow the infected cells to act as ‘Trojan horses’ for further dissemination to cause systematic infections^57^. Therefore, more effective elimination of intracellular bacteria has the potential to improve current antibiotic therapeutics.

Staphylococcal catalase shields intracellular *S. aureus* by degrading H_2_O_2_ generated by murine peritoneal macrophages^58^. Catalase has been reported to assist *Campylobacter jejuni* survive within macrophages as evidenced by extensive killing of catalase-deficient *Campylobacter jejuni* by macrophages^59^. Neutrophils, as the first line of defense, can also harbor pathogens inside the phagocytic vacuoles^60^, abetted by bacterial catalase that play a vital role in intracellular survival^61^. Therefore, we wondered whether photoinactivation of catalase could boost immune mechanisms of eliminating intracellular pathogens.

Thus, we infected the mice macrophage cell line RAW 264.7 with log-phased MRSA USA300 and 410 nm pre-treated log-phase MRSA USA300 at a multiplicity of infection (MOI) of 100 for one hour. After eliminating MRSA USA300 outside the macrophages by incubating with gentamicin for one hour, we adopted a protocol to label the live (SYTO 9)/dead (PI) MRSA USA300 inside the macrophages^62^. As shown in **Figure 6a-c**, there was a large number of live MRSA USA300 in the cytoplasm of macrophages after phagocytosis of untreated MRSA USA300. In comparison, the number of dead 410 nm pre-exposed MRSA USA300 is significantly increased inside macrophages (**Figure 6d-f**). We performed quantitative analysis of live/dead MRSA USA300 inside single macrophage from fifty macrophages (**Figure 6g-h**). There was a clear difference in the number of dead bacteria between the two groups, which indicates that 410 nm exposure reduces intracellular bacterial burden of macrophages.

**Figure 6.**
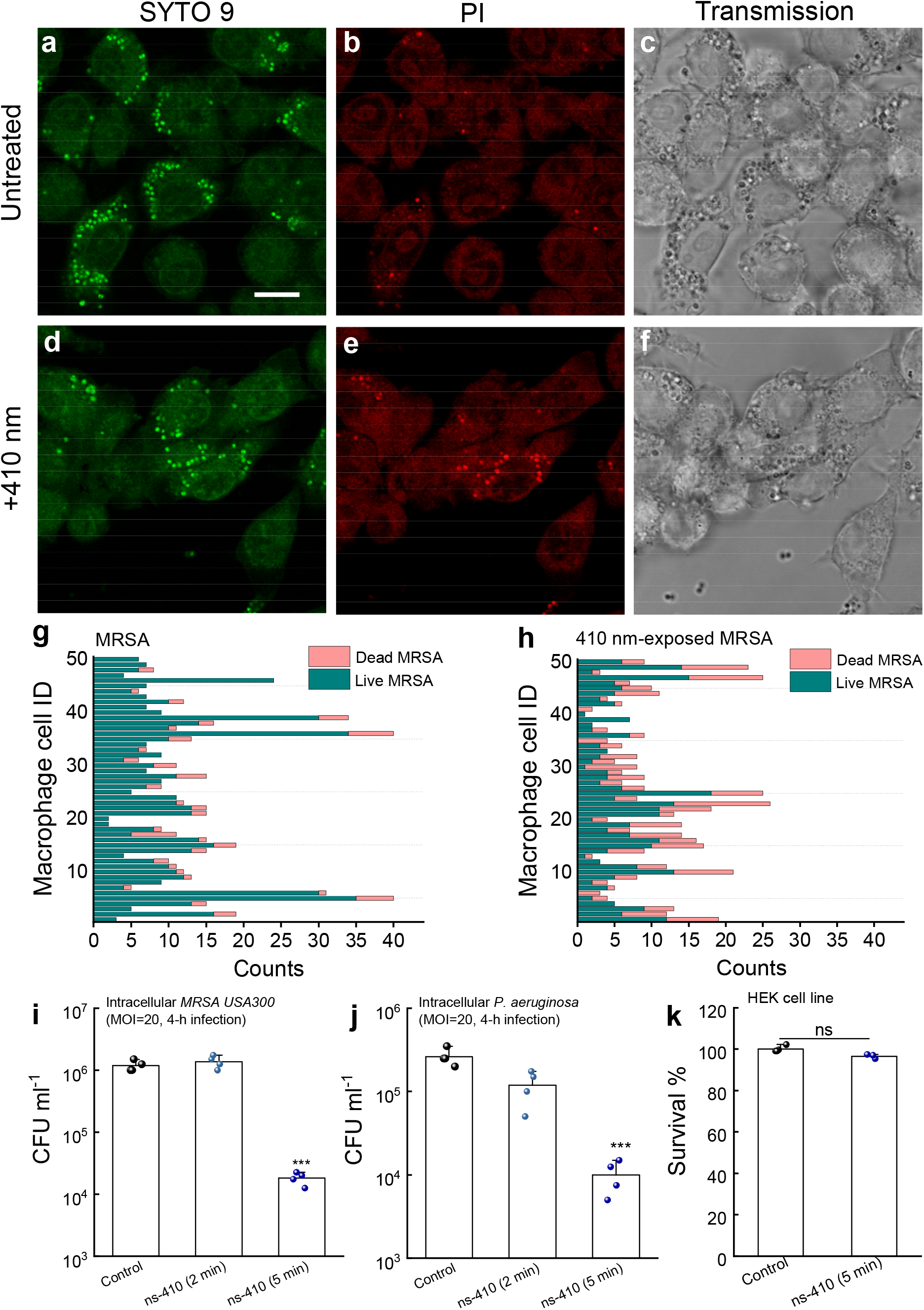
Photoinactivation of catalase assists macrophages to eliminate intracellular bacteria. **a**-**c**. Confocal images of live (SYTO 9, **a**), dead (PI, **b**) and corresponding transmission images (**c**) of intracellular MRSA USA300 inside RAW 264.7 macrophages after MRSA USA300 infected RAW 264.6 cells for 1 h at a multiplicity of infection (MOI) of 100 in serum-free DMEM media. **d**-**f**. Confocal images of live (SYTO 9, **d**), dead (PI, **e**) and corresponding transmission images (**f**) of intracellular MRSA USA300 inside RAW 264.7 macrophages after 410 nm-exposed MRSA USA300 infected RAW 264.7 cells for 1 h at a MOI of 100. 410 nm: 35 mW/cm^2^, 8 min. **g**-**h**. Quantitative analysis of the amount of live/dead MRSA inside single RAW 264.7 cells from the above two scenarios. **i**. CFU ml^-1^ of intracellular MRSA USA300 after MRSA (with/without 410 nm exposure) infected RAW 264.7 cells for 4 hours at a MOI of 20. **j**. CFU ml^-1^ of intracellular *P. aeruginosa* after *P. aeruginosa* PAO1 (with/without 410 nm exposure) infected RAW 264.7 cells for 4 hours at a MOI of 20. **k**. Survival percent of human epithelial keratinocyte (HEK) cell lines with/without 410 nm exposure. 410 nm: 50 mW/cm^2^. Data: Mean±SD. Student unpaired *t*-test. ***: *p*<0.001, **: *p*<0.01, ns: not significant.

To further confirm that photoinactivation of catalase enhances macrophage elimination of intracellular pathogens, we enumerated intracellular bacteria by lysing macrophages with 0.1% Triton-X for 3 min after 4-h infection. As shown in **Figure 6i**, there was an approximate 2-log_10_ reduction of intracellular MRSA USA300 burden in the 410 nm exposure (15 J/cm^2^) group. Consistently, a 1.5-log_10_ reduction of intracellular *P. aeruginosa* was obtained in macrophages infected with 410 nm (15 J/cm^2^)-exposed *P. aeruginosa* (**Figure 6j**). We showed that 410 nm exposure didn’t cause significant toxicity to a human epithelial keratinocyte (HEK) cell line (**Figure 6k**). Collectively, the results of the confocal live/dead imaging along with the CFU assay suggest that 410 nm exposure can facilitate host immune killing of intracellular pathogens.

### Photoinactivation of catalase reduces *P. aeruginosa* burden in a mouse skin abrasion model

Towards clinical translation, we further evaluated the synergy between photoinactivation of catalase and H_2_O_2_ using a bacterial infection murine model. Specifically, we adopted a *P. aeruginosa*-infected mouse skin abrasion model^63^ to mimic clinical *P. aeruginosa*-mediated skin infection. Briefly, we applied 10^8^ CFU of *P. aeruginosa* to abrazed mouse skin for three hours. After the wound was established, 410 nm blue light (120 J/cm^2^), 0.5% H_2_O_2_, 0.5% H_2_O_2_ plus 410 nm (120 J/cm^2^) were independently applied to the infected area for two times before euthanasia (**Figure 7a**). Wound tissues from the euthanized mice were then subjected to homogenization in order to enumerate CFU. As shown in **Figure 7b**, the untreated mice wound has around 10^6^ CFU/ml of *P. aeruginosa*, and 410 nm light alone significantly (*p*=0.02) reduced bacterial burden by around 60%. Noticeably, 410 nm light exposure significantly enhanced (*p*=0.0002) the bactericidal effect of 0.5% H_2_O_2_ against *P. aeruginosa* by one order of magnitude.

**Figure 7.**
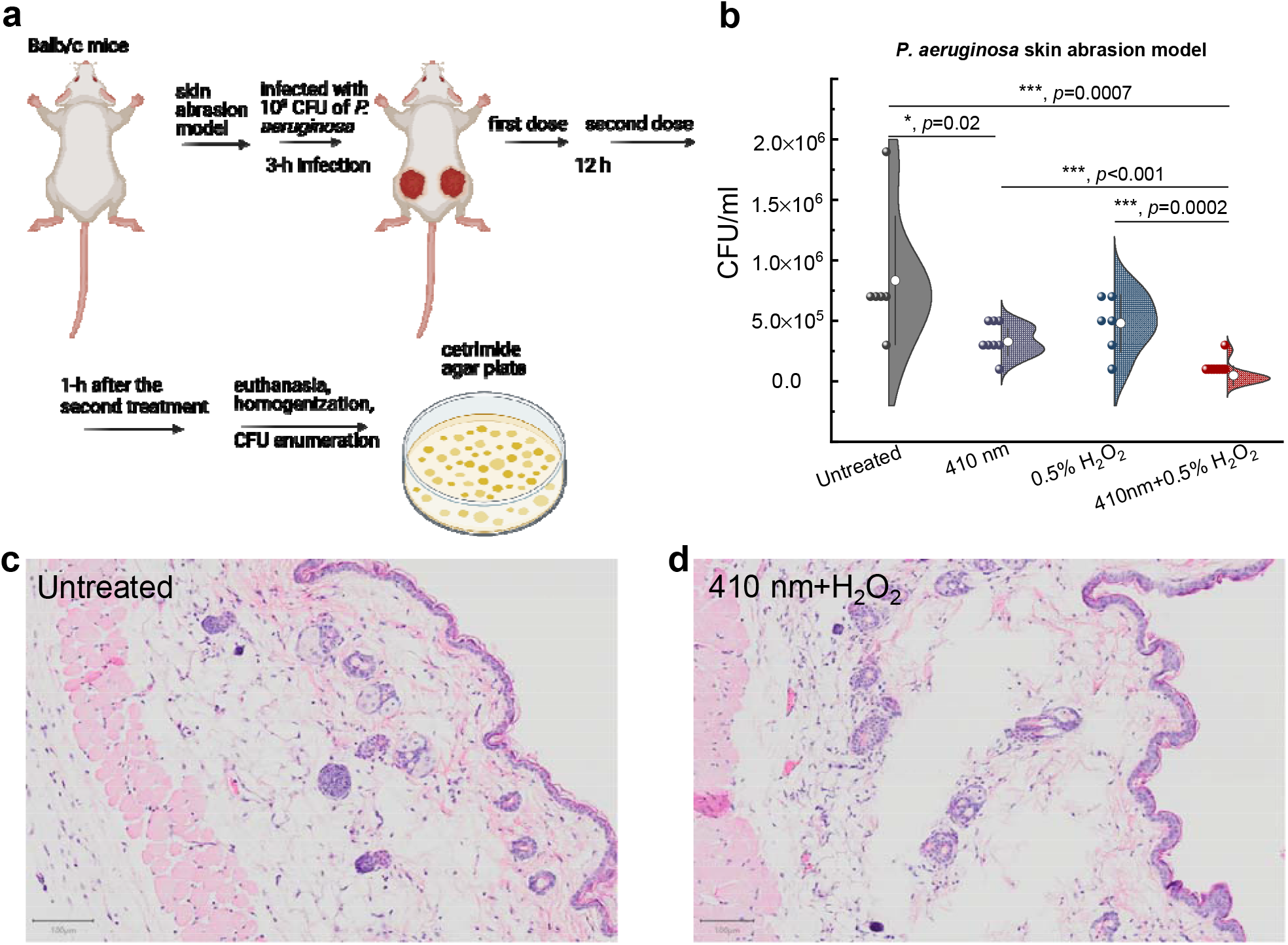
Photoinactivation of catalase reduces *P. aeruginosa* burden in a *P. aeruginosa*-induced mouse skin abrasion model. **a**. Schematic illustration of *in vivo* mice experiment. **b**. CFU ml^-1^ of *P. aeruginosa* PAO1 from the infected wound tissues among four different groups. **c**-**d**. Histology analysis of mice skin from untreated group along with 410 nm plus H_2_O_2_ treated group. Scalar bar=100 µm. Data: Mean±SD from at least six replicates. 410 nm: 120 J/cm^2^. H_2_O_2_: 0.5%. Student unpaired *t*-test. ***: *p*<0.001, **: *p*<0.01, *: *p*<0.05, ns: not significant. An outlier was removed based on the Dixon’s Q test and whisker box plot.

To evaluate whether our combinational treatment causes skin damage, we collected the mouse skin after applying 410 nm at the same dose plus 0.5% H_2_O_2_. We then performed the hematoxylin and eosin staining and histology analysis. As shown in **Figure 7c-d**, epidermis, dermis and subcutaneous tissues were not different from the untreated control. Collectively, these *in vivo* data provide the foundation for exploring the clinical utility of our catalase-targeted therapy against multidrug-resistant bacterial infections.

## Discussion

Since the discovery of penicillin in 1928 by Alexander Fleming^64^, there was a golden age (from 1940s to 1960s) for the discovery of antibiotics, most of which remain in clinical use today^65^. Yet, imprudent usage of antibiotics speeded up the natural selection of drug-resistant bacteria^66^. In the past decades, the ever-rising emergence of multidrug-resistant bacteria has been an alarming threat worldwide. Moreover, the pace of antibiotic development hasn’t kept up with antibiotic resistance development. Therefore, novel alternative approaches are highly desired to combat the new waves of multidrug-resistant bacterial infections.

It has been reported that catalase serves an important virulence function for intracellular bacteria evasion of neutrophil phagocytosis^67^. *E. coli* pretreated with phenazine methosulfate showed a nine-fold increase of catalase synthesis, and demonstrated resistance against neutrophil killing^68^. With catalase expression, *S. aureus* has significantly higher viability in the presence of human neutrophils^69^. Besides neutrophils, staphylococcal catalase protects intracellular bacteria by neutralizing H_2_O_2_ produced by macrophages^58^. Not only does catalase plays an important role in host-microbe interaction, it has been shown that catalase expression contributes significantly to the survival of catalase-positive *S. aureus* against catalase-negative *Streptococcus pneumoniae* in a murine model of nasal colonization^70^. Therefore, catalase inactivation could deprive pathogenic bacteria of an essential armament.

In this paper, we identified catalase as a key molecular target of blue light for a wide range of pathogens. Through spectroscopic study, we found that catalase can be functionally inactivated by blue light, especially at 410 nm. We further demonstrated that photoinactivation of catalase renders wide-ranging pathogenic bacteria and *P. aeruginosa* biofilms highly susceptible to subsequent H_2_O_2_ or H_2_O_2_-producing agents. The close correlation between catalase inactivation and its bacterial killing efficiency further validates that catalase is a major molecular target of blue light. In addition, photoinactivation of catalase significantly enhances macrophage killing of intracellular pathogens, and further reduces bacterial burden in a *P. aeruginosa*-infected mouse abrasion model without causing significant skin damage.

We found that ns-410 nm blue light is significantly more effective at inactivating catalase than CW-410 nm using the same dose. Moreover, ns-410 nm plus H_2_O_2_ kills more bacteria compared to CW-410 nm plus H_2_O_2_. Enhanced bacterial killing probably comes from the second-order photobleaching behavior of catalase under 410 nm irradiance, where prosthetic heme rings inside the tetramer might react with each other to detach from protein matrix. Raman spectra of catalase before and after 410 nm treatment hinted the porphyrin changes or dissociation. As an additional benefit, ns-410 nm exposure could also eliminate the heating issue which is always accompanied by CW-410 nm irradiance. It has been reported that heat dissipation is in the range of microsecond^71^, and ns-410 nm here was modulated at a frequency of 10 Hz, which could efficiently address the heat accumulation issue. In short, ns-410 nm might have better potential for clinical translation to treat multi-drug resistant bacterial infections compared to CW-410 nm.

We also found that stationary-phase pathogens such MRSA USA300 or *P. aeruginosa* are significantly more susceptible to subsequent H_2_O_2_ attack after photoinactivation of catalase. Stationary-phase or non-dividing pathogens are known to cause persistent infections such as endocarditis or osteomyelitis or biofilm-associated infections^72^. The persisting pathogens appear to be metabolically dormant and thrive under nutrition depleted environment^73^. The dormant bacteria demonstrate higher levels of tolerance and persistence to antimicrobial agents when compared to metabolically active planktonic bacteria^74^. Therefore, alternative approaches to combat stationary-phase pathogens is important. Catalase activity has been reported to be increased in the stationary-phase bacteria^75^. Our novel approach demonstrated bactericidal efficacy against a broad range of stationary-phase pathogens. This synergy effectively eliminated *P. aeruginosa* in a biofilm setting that is resistant to H_2_O_2_ alone.

We showed that photoinactivation of catalase also enhances immune cell elimination of intracellular pathogens. Bacterial catalase promotes intracellular pathogens survival inside immune cells^76^. As shown in **Figure 6a**, as significant portion of MRSA USA300 remained alive after internalization by macrophages. After photoinactivation of catalase, both intracellular MRSA USA300 and *P. aeruginosa* burden significantly reduced. Photoinactivation of catalase also significantly reduced bacterial load in the mouse skin abrasion model (**Figure 7b**) without causing skin damage. Although catalase is a major molecular target of blue light, it is not the only target of 410 nm irradiance since some H_2_O_2_ killing of catalase-deficient *E. coli* mutants and *S. aureus* mutants was still observed after 410 nm blue light treatment (**Figure 2**). This finding indicates that molecular targets of the 410 nm light other than catalase likely exist. Noteworthy, there are other endogenous molecules which are intrinsic to specific bacteria already revealed as molecular targets of blue light. Collectively, our findings reveal a significant mechanism of antimicrobial blue light and fueled a novel catalase-targeted strategy to combat clinical multidrug-resistant bacterial infections.

## Methods and Materials

### Blue light source

Pulsed blue light was administered using a Opolette HE 355 LD laser (OPOTEK). The nanosecond pulsed laser is tunable from 410 nm to 2200 nm, has a repetition rate of 10-20 Hz, and a pulse width of ∼5-10 ns. Using a collimator attached to an optical fiber, the diameter of the laser beam was expanded to 10 mm. With these set parameters, the pulsed laser provides a power output ranging from 25 to 100 mW/cm^2^, depending on the laser wavelength used. Continuous wave blue light was delivered through a mounted 405 nm blue light LED (M405L4, Thorlabs) with an adjustable collimation adapter (SM2F32-A, Thorlabs) focusing the illumination region to a ∼1 cm^2^ region. A T-Cube LED driver (LEDD1B, Thorlabs) allowed for adjustable light fluencies up to 400 mW/cm^2^.

### Bacterial strains and cell lines

Bacterial strains MRSA (USA300), *P. aeruginosa* PAO1 (ATCC 47085), *P. aeruginosa* 2 (ATCC 1133), *P. aeruginosa* 3 (ATCC 15442), *P. aeruginosa* 4 (ATCC 9027), *S. enterica* 2 (ATCC 700720), *S. enterica* 3 (ATCC 13076), *E. coli* 1 (BW25113), *K. pneumoniae* 1 (ATCC BAA 1706), *A. baumannii* 1 (ATCC BAA 1605), *A. baumannii* 2 (ATCC BAA-747), *E. faecalis* 1 (NR-31970), and *E. faecalis* 2 (HM-335) were provided by the Dr. Mohamed N. Seleem at Virginia Tech. *E. coli* mutants (*ΔahpC, ΔkatG, ΔkatE, ΔkatGE*) were provided by the Dr. Xilin Zhao Group at Rutgers University. All cell lines used in this study, including the RAW 264.7 murine macrophages, chinese hamster ovary (CHO) cells, and human embryonic kidney-293 (HEK) cells, were purchased directly from the American Type Culture Collection (ATCC).

### Transient absorption imaging of real-time photobleaching of catalase

As described previously^77^, an optical parametric oscillator synchronously pumped by a femtosecond pulsed laser generated the pump (820 nm) and probe (1040 nm) pulse trains. The pump and probe beams were then frequency-doubled via the second–harmonic generation process to 410 nm and 520 nm through barium borate crystals, respectively. Temporal delay between the pump and probe pulses was controlled through a motorized delay stage. The pump beam intensity was modulated with an acousto-optic modulator. The intensity of each beam was adjustable through the combination of a half-wave plate and a polarization beam splitter. Thereafter, pump and probe beams were collinearly combined and directed into a laboratory-built laser scanning microscope. Through the nonlinear process in the sample, the modulation of pump beam was transferred to the unmodulated probe beam. Computer-controlled scanning galvo mirrors were used to scan the combined laser beams in a raster scanning approach to create microscopic images. The transmitted light was collected by an oil condenser. Subsequently, the pump beam was spectrally filtered by an optical filter, and the transmitted probe intensity was detected by a photodiode. A phase-sensitive lock-in amplifier (Zurich Instruments) then demodulated the detected signal. Therefore, pump-induced transmission changes in the probe beam versus the temporal delay can be measured. This change over time delay shows different time-domain signatures of a chromophore, thus offering the origin of the chemical contrast.

### Raman spectra of catalase before and after 410 nm irradiance

Raman spectra of catalase before and after 410 nm exposure were acquired under a Horiba Raman system (1221, LABRAM HR EVO, Horiba) with a 40× objective under the excitation of 532 nm. Raman spectra were obtained under a 20-s acquisition time with a 10% laser ND filter. Bovine liver catalase (C9322, Sigma Aldrich) was dissolved in sterile distilled H_2_O (4 U/mL) and then air-dried onto a clean cover slide. Changes in catalase structure were evaluated by examining decreases in specific Raman peaks following light treatment.

### Quantitation of remaining active catalase

Measurement of active catalase was primarily quantified through the use of an Amplex Red Catalase Assay (A22180, Thermo Fisher Scientific). Briefly, solutions containing catalase (either bovine liver catalase or catalase-positive bacteria inoculum) were treated with blue light, after which 25 µL of the light treated solution was incubated with 40 µM of H_2_O_2_ for 30 minutes at room temperature within a 96-well plate. Following H_2_O_2_ treatment, 50 µL of a reaction stock containing 100 µM of Amplex Red and 0.4 U/mL of horseradish peroxidase were added to each well and the combination was incubated for 30 minutes at 37°C. Following incubation, the fluorescence of each well was measured using an excitation wavelength of 543 nm and an emission wavelength of 585 nm. In addition to the light treated samples, a phosphate buffered saline (PBS) negative control and an untreated positive control also treated with the assay in order to determine the remaining catalase percentage. Active catalase percent was calculated through the following equation:

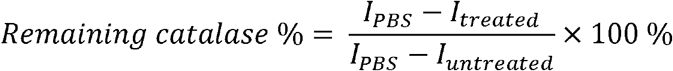

### CFU enumeration assay to evaluate the synergy between photoinactivation of catalase and ROS-generating agents

*P. aeruginosa* and MRSA were cultured overnight in TSB at 37°C within a shaking incubator (250rpm). The next day, the bacteria were suspended in 1×PBS (OD_600_ = 1.0), after which a 10 µL of aliquot was placed on a glass cover slide and then treated with ns-410 nm or CW-410 nm. Following light treatment, the droplet was transferred to a tube containing either PBS or 22 mM of H_2_O_2_ diluted in PBS. Samples were incubated for 30 minutes under 37°C, after which the bacteria samples were 10-fold serially diluted within a 96-well plate, plated on tryptic soy agar plates overnight, and then CFU enumerated the next day. Subsequent experiments testing the synergy between ns-410 or CW-410 blue light and different concentrations of H_2_O_2_ were also performed on other stationary phase bacteria including *E. coli, P. aeruginosa*, MRSA, *A. baumannii, S. enterica, K. pneumoniae*, and *E. Faecalis*. Similar to previous CFU synergy experiments, wild type *E. coli* (BW25113), an alkyl hydroperoxide reductase negative mutant strain (*ΔahpC*), and three catalase negative mutants (*ΔkatG, ΔkatE, ΔkatGE*) were all cultured overnight in TSB at 37°C within a shaking incubator (250rpm). Following incubation, bacteria strains were suspended in 1×PBS (OD_600_ = 1.0). A 20 µL aliquot of bacteria was placed on a glass coverslip and exposed to ns-410 light (32 mW/cm^2^, 14 J/cm^2^). The droplet was then removed and diluted with 780 µL of PBS. This light treated sample, alongside a non-light exposed sample, was treated with 2.2 mM of H_2_O_2_. Samples were shaken and incubated at 37°C for up to 2 hours. During the incubation, 60 µL aliquots would be removed from each sample tube for serial dilatation and CFU plating at the 20 minute, 40 minute, 1 hour, 1.5 hour, and 2-hour time points.

### Mammalian cell toxicity assay

To evaluate the potential toxicity of 410 nm exposure and short term, high concentration H_2_O_2_ exposure against mammalian cells, a MTT assay was performed using human epithelial keratinocyte (HEK) cell line and Chinese Hamster Ovary (CHO) cell line. HEK/CHO cells were cultured in Dulbecco’s Modified Eagle Medium (DMEM, Gibco) alongside 10% fetal bovine serum (FBS) until 90% confluence was achieved. Once high confluence was established, cells were removed through trypsin treatment, quantified with a cell counter, and diluted in DMEM media to a final cell concentration of 1×10^6^ cells/mL. 100 µL of cell media were added to each well of a 96-well plate, providing each well with 1×10^5^ cells per well. Each well was supplemented with an additional 100 µL of DMEM media to bring the final volume of each well to 200 µL. HEK/CHO cells were then incubated overnight at 37°C with 5% CO_2_ in order to allow the cells to adhere to the wells. Following the replacement of DMEM with PBS, light treated wells were exposed to 75 J/cm^2^ of 410 nm LED light (250 mW/cm^2^). After light exposure, PBS was removed and replaced with DMEM. For the H_2_O_2_ treatment groups, 0.1% H_2_O_2_ suspended in DMEM was added to the H_2_O_2_ treatment groups for 1 minute, after which the H_2_O_2_ containing media was removed and the wells were washed twice with PBS. Identical two-fold PBS washing was also applied to the other treatment groups to maintain experimental consistency. After that, the PBS was replaced with fresh DMEM and incubated overnight at 37 C with 5% CO_2_ to allow for the surviving cells to recover from the stress exerted by the treatment. The next day, an MTT viability assay was performed based on previously established protocols. MTT absorbance measurements were quantified via plate reader at 590 nm. The assay was performed in replicates of N=4.

### CFU assay between photoinactivation of catalase and certain antibiotics

For each antibiotic tested, bacterial strains were usually pre-cultured in antibiotics prior to light exposure of subsequent incubation. To summarize, 1 mL of overnight cultured bacteria was centrifuged and suspended in 1 mL of fresh TSB media supplemented with either 10 µg/mL of tobramycin (Sigma Aldrich, T4014) or 0.1 µg/mL of ciprofloxacin (Sigma Aldrich, 17850). Following pre-culture treatment, the bacterial solution was spun down and resuspended in 1×PBS. A 10 µL aliquot of the bacterial preculture stock was then placed on a coverslip and exposed to ns-410 with a dose of 18 J/cm^2^, after which the aliquot was transferred to 990 µL of fresh TSB supplemented with either 2 µg/mL of tobramycin or 0.1 µg/mL of ciprofloxacin and incubated at 37°C for up to 6 hours. CFU dilution and enumeration was taken at various time points during the incubation period.

### Intracellular bacteria assay

In all experiments, bacteria were cultured in TSB. To assess intracellular killing with macrophage RAW 264.7 cell line, MRSA USA300 or *P. aeruginosa* was taken from an exponentially growing culture and washed in PBS. Macrophages were pre-washed with serum-free DMEM media immediately before infection, and infected by MRSA USA300/P. *aeruginosa* with and without 410 nm treatment. Then co-culture was incubated at 37 °C in a humidified tissue culture incubator with 5% CO_2_ to permit phagocytosis of the bacteria. After 2-4 h, the infection mix was removed and replaced with normal growth media (DMEM supplemented with 10% FBS, 10 mM HEPES) and gentamycin was added at 50 μg/ml to prevent growth of extracellular bacteria for 1 hour. Two approaches were utilized to evaluate the difference. The first one was to utilize a Live/Dead confocal staining assay to visualize the live and dead bacteria inside macrophages, respectively. Briefly, after fixation with 10% formalin following gentamicin treatment, samples were permeabilized with 0.1% Triton-X for 3 min at r.t. After that, a Live/Dead fluorescence kit (Thermo Fisher Scientific, L7007) was utilized to stain the intracellular bacteria. Confocal laser scanning microscope (FV3000, Olympus) was employed to visualize stained samples. The second one was to enumerate the intracellular CFU. In brief, co-cultures were permeabilized with 0.1% Triton-X for 3-5 min, vigorous pipetting was conducted to release intracellular bacteria. 10-fold serial dilution was immediately applied to count the viable intracellular bacteria.

### *P. aeruginosa* biofilm assay

As reported previously^10^, *P. aeruginosa* biofilms were formed onto a coupon through a CDC biofilm reactor for two days. After the biofilm formation, different treatment schemes were then applied. Live/dead fluorescence staining was utilized to evaluate the treatment efficacy. Confocal laser scanning microscope was then employed to capture the counterstained images.

### *In vivo* murine infection model and histology

Twenty BALB/c mice (Jackson Laboratories, 000651) were placed under anesthesia and a #15 sterile scalpel was used to generate a 1 cm^2^ abrasion wound by carefully scraping the epidermis of the skin without drawing blood. Once the wound had been generated, a 10 µL aliquot containing 10^8^ CFU of log-phase PAO1 in PBS was placed onto the abrasion wound and spread evenly across the wound with a pipette tip. Once the droplet had dried, the mice were returned to their cage to allow the mice to recover from their abrasion wound and to provide time for the bacteria to infect the wound. The 20 mice were then divided into four treatment groups, each consisting of 5 mice (N = 5): Untreated, 410 nm treated, H_2_O_2_ treated, and 410 nm plus H_2_O_2_ treated. Light treatment was applied by positioning mice under 200 mW/cm^2^ (ANSI standard) 410 nm LEDs and exposing their bacteria infected wounds to 120 J/cm^2^ of blue light. For H_2_O_2_ treatment, 10 µl of 0.5% H_2_O_2_ was evenly distributed on the infected wounds and allowed to naturally dry. Combination treatment consisted of the application of previously described light treatment followed by H_2_O_2_ treatment. Treatments were applied to mice twice over the course of 15 hours, with the first treatment being applied 3 hours following infection and the second treatment being applied 12 hours after the first treatment. 3 hours after the second treatment, mice were euthanized and wound tissue was harvested, homogenized, and serially diluted. CFU enumeration was performed on *P. aeruginosa*-specific cetrimide agar plates (Sigma Aldrich, 22470).

The potential phototoxicity of the treatment on the skin was evaluated by applying the combination treatment on healthy, non-wounded skin regions present on the combination treated mice. This healthy region of the skin would receive the same two treatments as the wound site, and during tissue harvesting, this region was excised and preserved in 10% buffered formalin alongside a collection of unwounded skin samples from the untreated mice. Formalin fixed samples were submitted to the Boston University Experimental Pathology Laboratory Service Core for histology processing and hematoxylin and eosin (H&E) staining. Histology slides were then visualized under an inverted microscope through a 60× objective.

### Statistical analysis

Statistical analysis was conducted through student unpaired *t*-test and One-way ANOVA. *** means significantly different with a *p*-value < 0.001. ** means significantly different with a *p*-value < 0.01. * means significantly different with a *p*-value < 0.05. ns means no significance.

## Acknowledgments

This work was partly supported by R01AI141439 to J.-X.C. We kindly acknowledge Dr. Xilin Zhao from Rutgers University for providing the catalase-deficient *E. coli* strains. Research reported in this publication was also supported by the Boston University Micro and Nano Imaging Facility and the office of the director, National Institute of Health, National Institute of Health under Award Number S10OD024993.

## Author contributions

P.-T.D. and J.-X.C. conceived the synergistic therapeutic treatment between photoinactivation of catalase and H_2_O_2_ or certain antibiotics. P.-T.D., Y.F.Z., J.H. discovered that catalase from catalase-positive bacteria could be ubiquitously inactivated by 410 nm. P.-T.D. characterized catalase photoinactivation and intracellular bacteria assay. P.-T.D., S.J. and J.H. conducted the *in vitro* CFU assay. J.H. conducted the biofilms experiment. P.-T.D., S.J. and Y.W.Z. conducted the *in vivo* mice abrasion experiments and histology assay. P.-T.D. and J.-X.C. co-wrote the manuscript. G.L. provided constructive suggestions over the project and manuscript. All authors read and commented on the manuscript.

## Supplementary information for manuscript

**Supplementary Figure 1.**
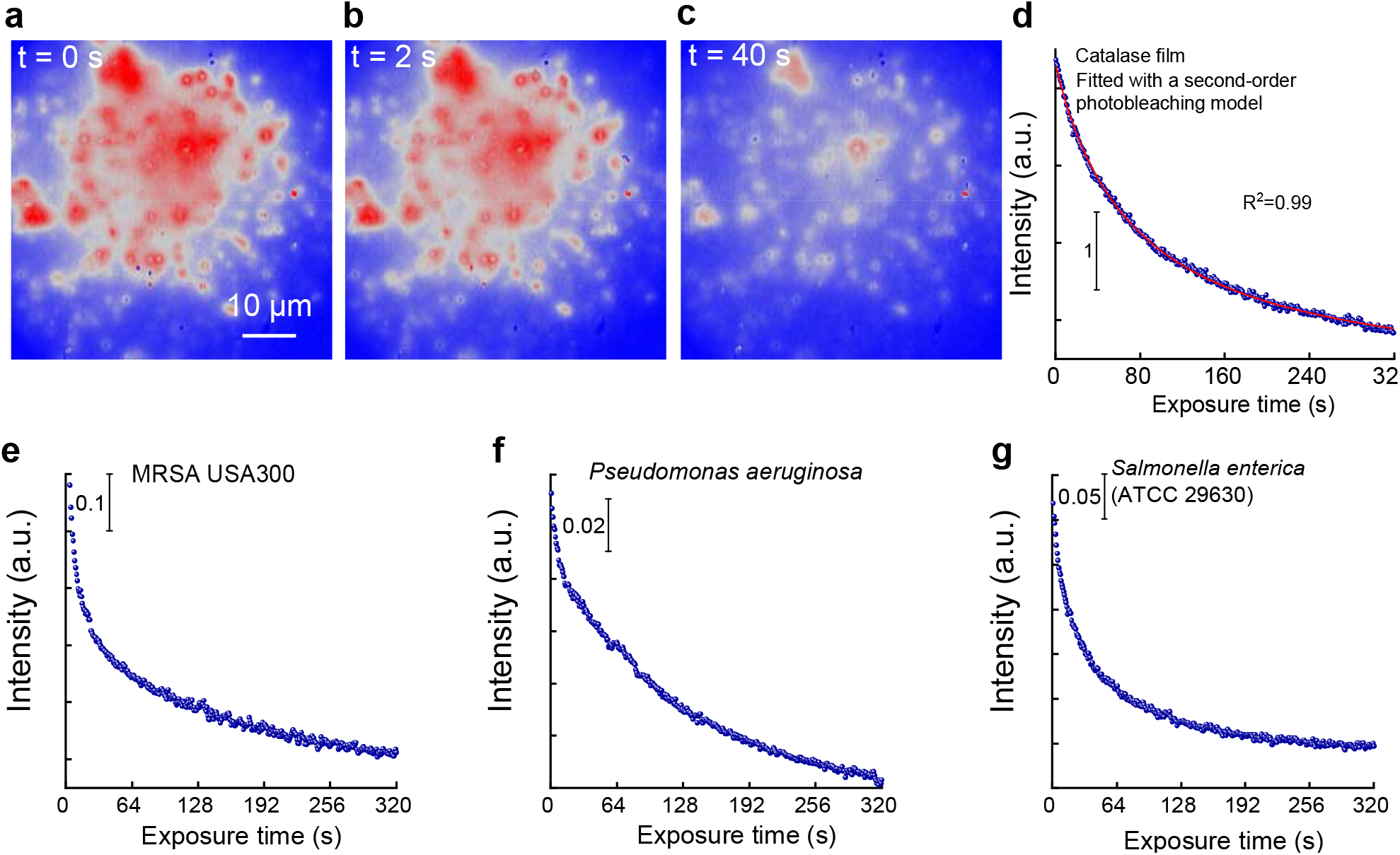
Characterization of photoinactivation of catalase under transient absorption microscope. **a**-**c**. Transient absorption images of bovine liver catalase dried on a cover slide under different exposure time. **d**. Time-lapse transient absorption signals of dried bovine liver catalase. Curve fitted by a second-order photobleaching model. **e**-**g**. Time-lapse transient absorption signals of dried MRSA USA300 (**e**), *P. aeruginosa* (**f**), and *Salmonella enterica* (**g**). Pump=410 nm, 5 mW on the sample; probe=520 nm, 7 mW on the sample. Scalar bar=10 µm.

**Supplementary Figure 2.**
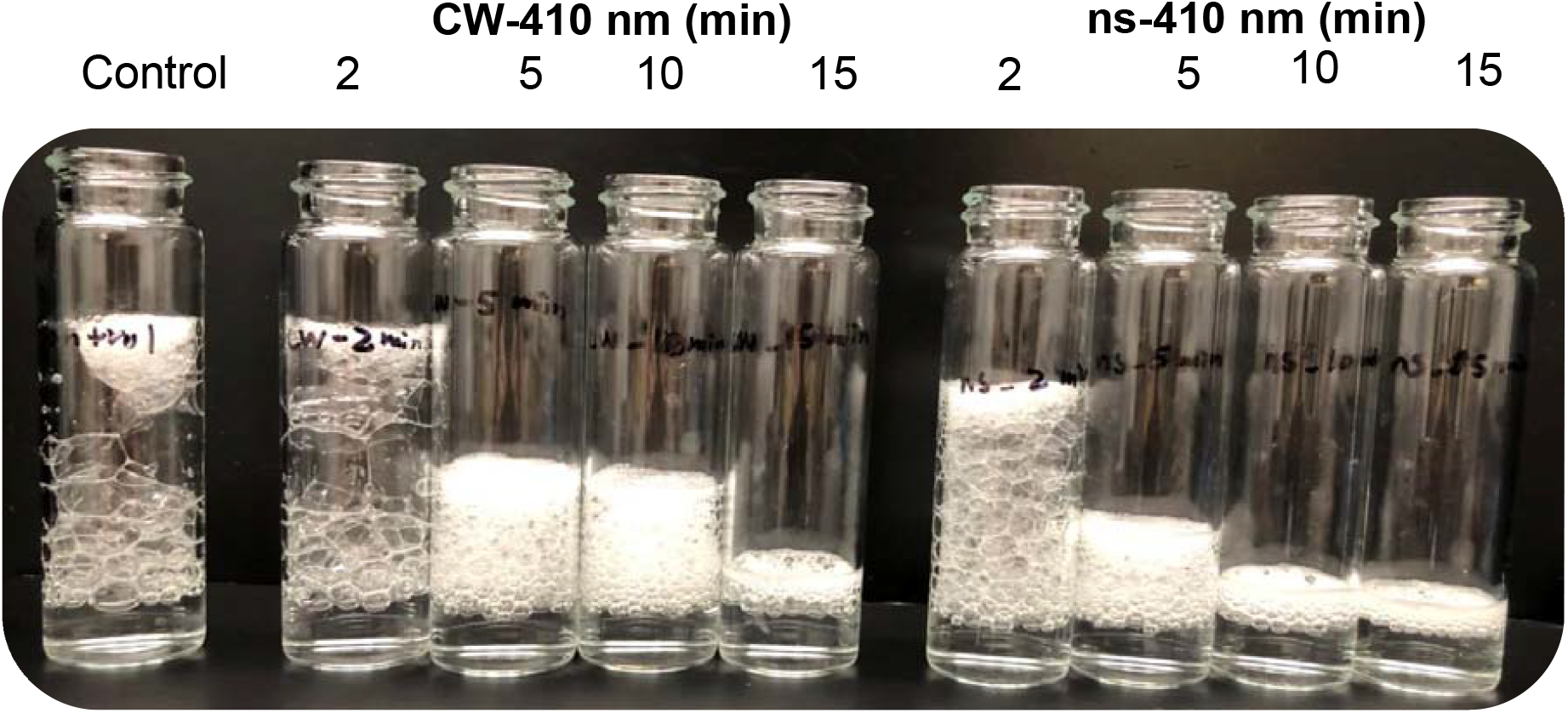
Bubble test comparison between CW-410 nm and ns-410 nm exposure on the capability to inactivate bovine liver catalase. 410 nm: 50 mW/cm^2^.

**Supplementary Figure 3.**
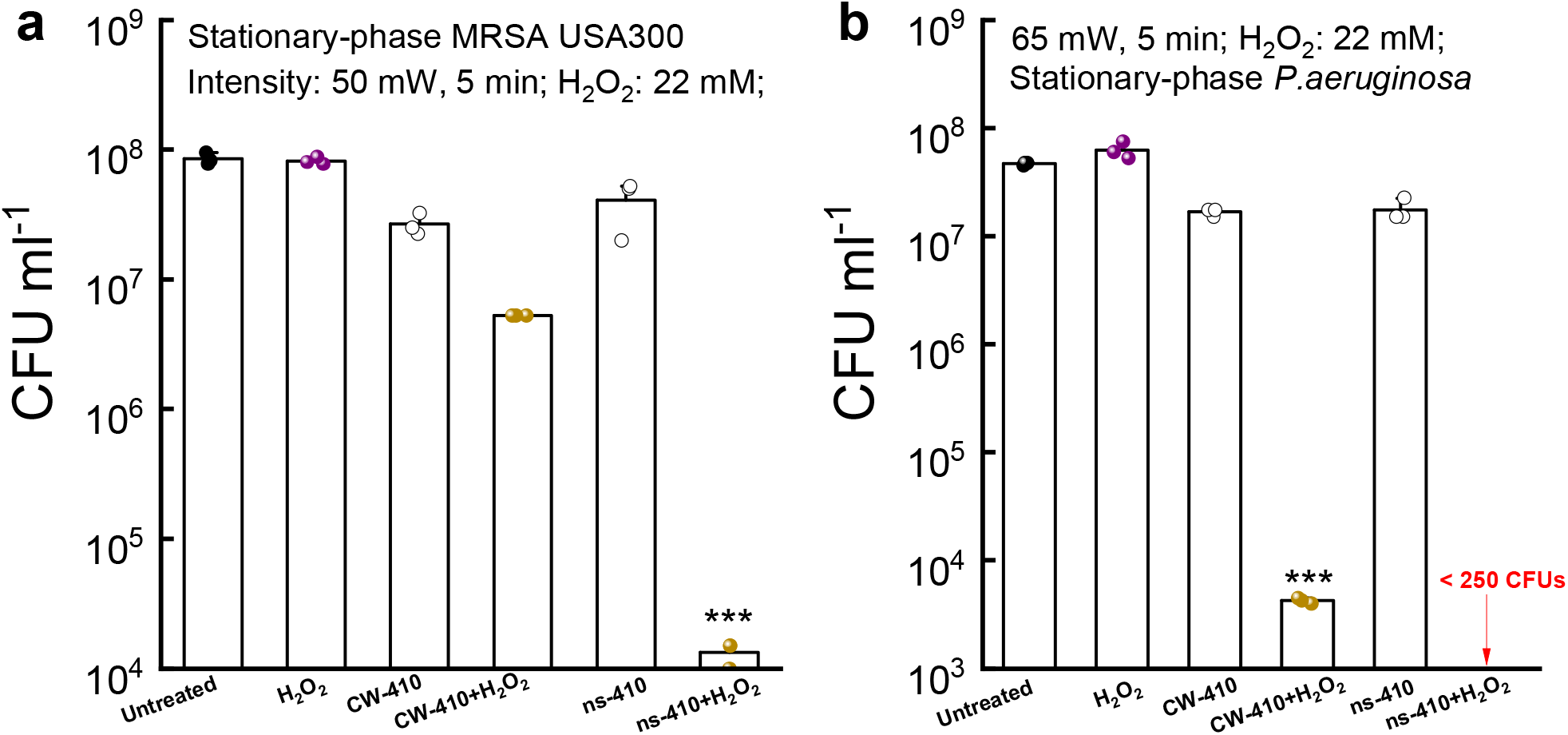
CFU ml^-1^ of stationary-phase MRSA USA300 (a) and *P. aeruginosa* PAO1 (b) under CW-410 nm and ns-410 nm treatments. H_2_O_2_: 30-min incubation time. Data: Mean±SD. Student unpaired *t*-test. ***: *p*<0.001.

**Supplementary Figure 4.**
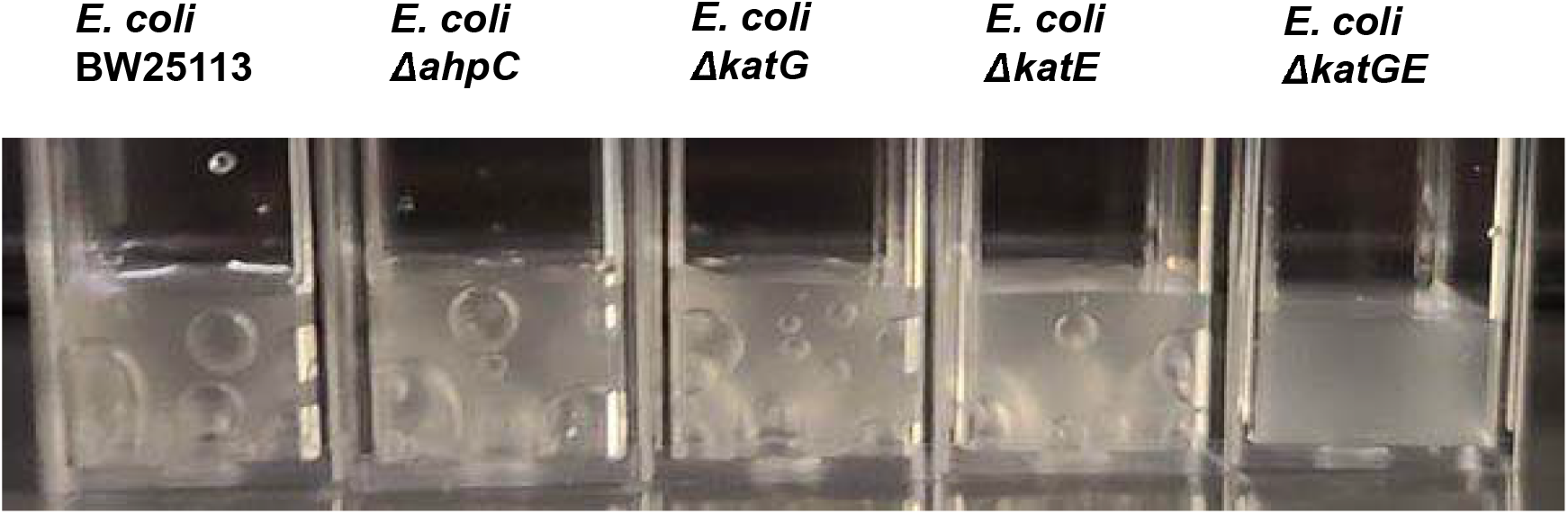
Bubble formation of different *E. coli* strains in the presence of 3% H_2_O_2_.

